# blkbox: Integration of multiple machine learning approaches to identify disease biomarkers

**DOI:** 10.1101/123430

**Authors:** Boris Guennewig, Zachary Davies, Mark Pinese, Antony A Cooper

**Affiliations:** Garvan Institute of Medical Research, Darlinghurst, New South Wales, Australia; St Vincent’s Clinical School BABS UNSW Australia, Sydney, New South Wales, Australia; BABS UNSW Australia, Sydney, New South Wales, Australia

**Author notes:** Contributed equally to this work (Boris Guennewig, Zachary Davies).

## Abstract

**Motivation:** Machine learning (ML) is a powerful tool to create supervised models that can distinguish between classes and facilitate biomarker selection in high-dimensional datasets, including RNA Sequencing (RNA-Seq). However, it is variable as to which is the best performing ML algorithm(s) for a specific dataset, and identifying the optimal match is time consuming. blkbox is a software package including a shiny frontend, that integrates nine ML algorithms to select the best performing classifier for a specific dataset. blkbox accepts a simple abundance matrix as input, includes extensive visualization, and also provides an easy to use feature selection step to enable convenient and rapid potential biomarker selection, all without requiring parameter optimization.

**Results:** Feature selection makes blkbox computationally inexpensive while multi-functionality, including nested cross-fold validation (NCV), ensures robust results. blkbox identified algorithms that outperformed prior published ML results. Applying NCV identifies features, which are utilized to gain high accuracy.

**Availability:** The software is available as a CRAN R package and as a developer version with extended functionality on github (https://github.com/gboris/blkbox).

## 1 Introduction

Transcript expression is the most informative high throughput modality for predicting clinical phenotype (Ray et al., 2014). Transcriptomic data is high dimensional, with very large numbers of transcripts (features or p > 10,000), but a comparatively small number of biological replicates (n = 2 to 1000). Although transcriptional data can be highly predictive of biological state, extracting robust and informative insights from such complexity is a longstanding challenge in data analysis (Capobianco, 2014). Machine Learning (ML) algorithms are statistical methods that leverage this data complexity through ‘learning’ from data (James *et al*., 2013). By identifying highly predictive features, ML can identify transcripts useful to robustly diagnose patients or identify pre-symptomatic individuals (Iizuka et al., 2003; Klöppel et al., 2009). Unfortunately, ML methods are computationally demanding and currently underutilized (Libbrecht and Noble, 2015) due to the large number of known human transcripts. The computational requirements of ML methods can be reduced by restricting analysis to only the most informative transcripts, with little loss of accuracy (Fig. 1) (Xing *et al*., 2001). However, the process of selecting informative transcripts, termed feature selection, is complex: no single best method exists, and it is known that optimal input features can differ between ML algorithms (Suppl. Fig. 1b). This diversity explains the varying performance of ML algorithms on different input data. In machine learning the biggest challenge is selecting the best algorithm (James et al., 2013), which highlights the necessity to test multiple algorithms to obtain the most accurate model for a specific dataset. This need to explore multiple algorithms can be time-consuming, computationally challenging and hinders the application of ML approaches (Ray et al., 2014). Here we describe blkbox, a tool to simplify the exploration of ML algorithms. blkbox is simple to use (includes a shiny interface; (Suppl. Fig. 2).), provides comparative output over nine relevant ML algorithms, provides enhanced efficiency in potential biomarker selection through scored feature lists, reduces computational cost through optional feature selection, uses accepted standards in the ML field, is under constant development, and provides publication ready graphical results to the user. With a consistent interface, blkbox provides multiple high-performance ML algorithms to users without expertise in the ML field, streamlining access to these methods for researchers.

**Fig. 1.**
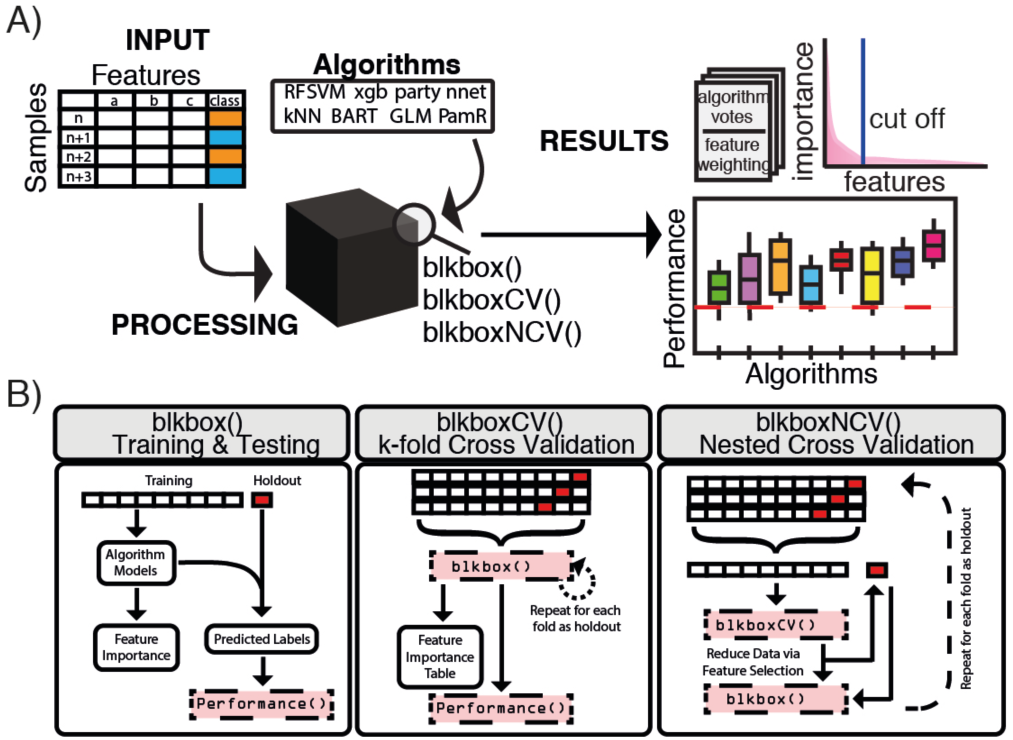
A): Schematic workflow and pipeline of blkbox. B) Schematics of the three model types implemented in blkbox.

## 2 Workflow

blkbox implements nine representative high-performance ML algorithms (Fig. 1A and Suppl. Methods) as well as functions for a range of ML workflows including standard training and testing (blkbox), cross fold validation (blkboxCV), and nested cross fold validation (blkboxNCV) (Fig. 1B). blkbox provides the functionality to plot different metrics (e.g. AUC), allowing comparison of algorithm effectiveness for a given dataset (Suppl. Fig. 3 & 4). The intersection of selected features in each testing holdout can be visualized in a Venn diagram or heatmap to identify commonly used features and assess variance among holdouts (Supp. Fig. 5). Importance of each feature can be found for each fold, tables present the values weighted by their performance on holdouts, enabling feature exploration and additional calculations of feature stability across holdouts (Vignette: blkbox).

## 3 Performance

blkbox was benchmarked against a published cancer transcription dataset (Suppl. Fig. 3) of 7070 features to determine running speed and accuracy. blkbox identified a combination of feature selection and machine learning algorithms (PamR-GLM) that outperformed the published results (red line in Suppl. Fig. 3). The identified combinations can be a starting point for further parameter optimization to achieve increased accuracy and/or robustness. Extraction of well performing features subsets provides both potential biomarker discovery and insights into underlying biological mechanisms. blkbox can operate without feature selection/reduction (Fig. 1A/B) but, based on the high computational resources required (Suppl. Fig. 3), this may only be possible on smaller datasets, under exclusion of cross fold validation, or without using resource hungry algorithms (Suppl. Fig. 3). Due to these constraints and the large size of most RNA-Seq datasets, a feature reduction step can be used in blkbox based on the rationale that not every feature contributes an importance to the statistical model and some can therefore be discarded. The cumulative sum of each feature’s relative importance (Fig. 1A upper right) provides an approximate area under the curve (AUC). Multiplication of the AUC by an AUC cutoff (0.0 – 1.0) can determine a threshold; features with a relative importance below the threshold are removed, and those retained are referred to as surviving features and used in the subsequent analysis.

## 4 Applications

blkbox is designed for transcriptomics analysis and exploration for biomarkers although its application can be applied to any question with a binary outcome. The heavily weighted transcripts identified by blkbox identify biomarker candidates to be tested, individually or in combination, for their capacity to robustly distinguish healthy individuals from patients or to identify pre-symptomatic patients. Analysis of the biological function of these transcripts can additionally provide mechanistic insights into the basis of the disease that may in turn lead to new therapies.

## 5 Conclusions

blkbox provides an intuitive and easy to use interface for the exploration of numerous ML algorithms. Performance metrics on each algorithm are presented in a clear and publication ready format and allows identification of top performing combinations. Feature lists of importance in accurate classification models enables the extraction of feature subsets, which in turn are promising biomarker candidates as well as providing potential insights into mechanisms underlying disease. The three functions (train & test, cross-fold validation, and nested cross fold validation) address pitfalls including over fitting and cope with real world heterogeneity. blkbox provides both an entry level capacity for new ML users due to its clear and simple syntax as well as providing advanced running scenarios. Feature selection methods reduces computational cost and enable head-to-head comparisons of algorithms, which would otherwise be impossible due to their exponential computational cost with large p.

The combination of those functions makes blkbox unique in the ML landscape and promises to find broad application in multiple scenarios beyond those discussed here.

## 6 Acknowledgements

The authors are thankful to Peter Holberton for a code review and to John S Mattick, Eugene Dubossarsky, Timothy J Peters and Max Skipper for helpful discussions.

## 7 Funding

This work was supported in part by grants from the NHMRC (APP1067350) and a Geoff and Dawn Dixon Fellowship to B. G. and A. A. C. We gratefully acknowledge the Swiss National Science Foundation for the funding of B. G. (P2EZP3_152143).

## Conflict of Interest

none declared.

---

**Suppl. Figure 1:**
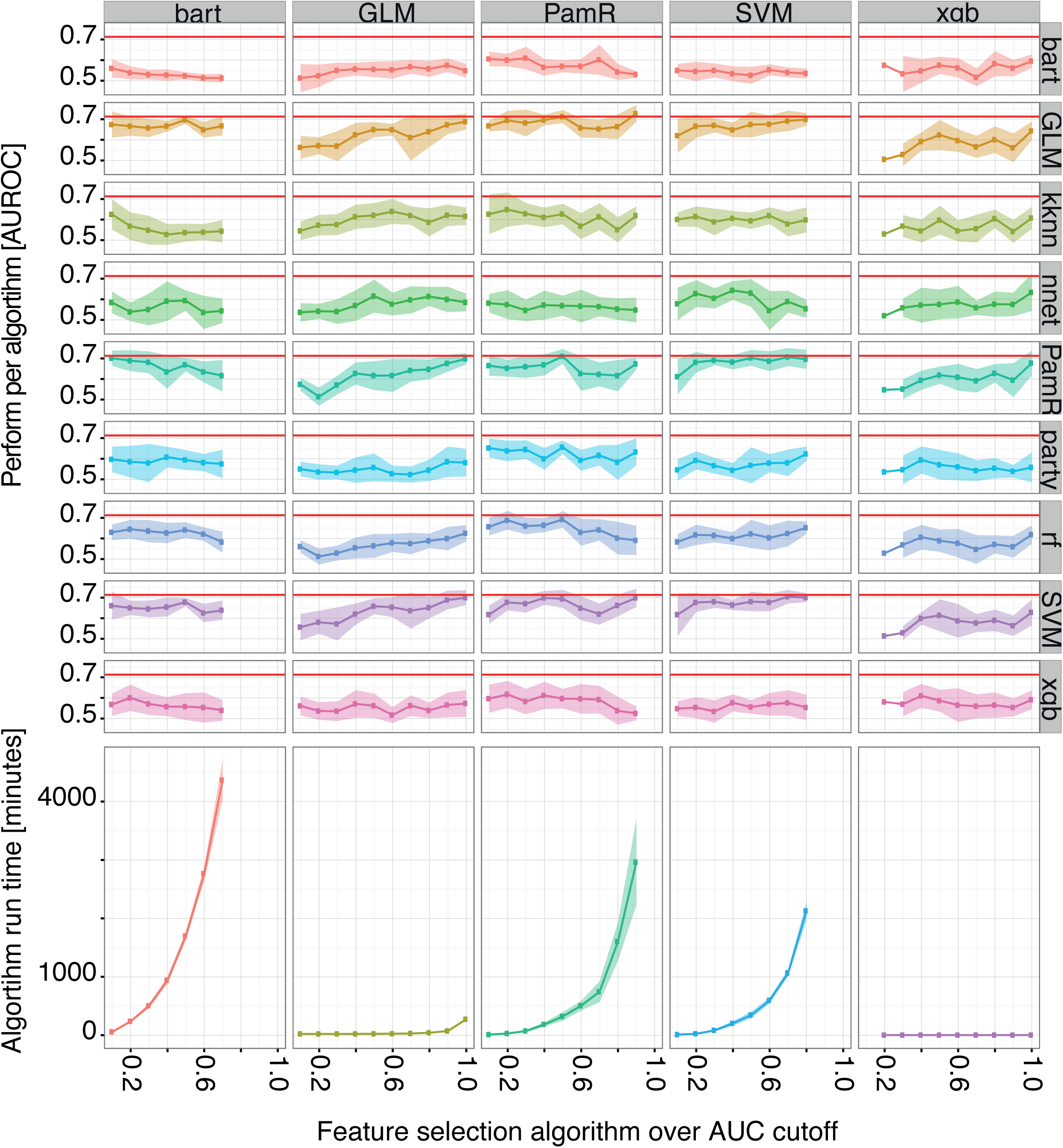
Exemplary benchmark on a 7070 features gene expression set (Iizuka et al., 2003), using 5/8 implemented features selection methods (columns) over AUC thresholds 0.1-1 and determining the AUROC with the included algorithms (rows) against the published result (red line from Statnikov et al., 2008). Iizuka, N. et al. (2003) Oligonucleotide microarray for prediction of early intrahepatic recurrence of hepatocellular carcinoma after curative resection. The Lancet, 361, 923–929 Statnikov, A. et al. (2008) A comprehensive comparison of random forests and support vector machines for microarray-based cancer classification. BMC Bioinformatics, 9, 319.

**Suppl. Figure 2:**
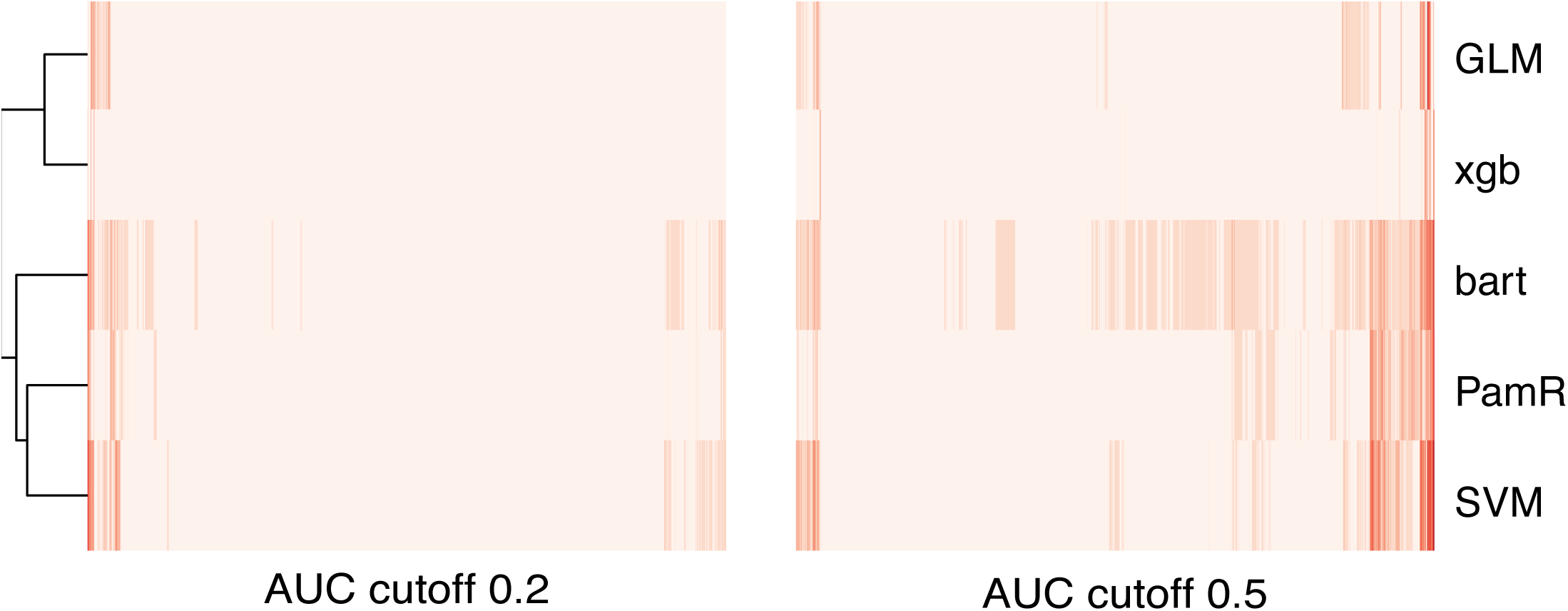
Partial heat maps of feature importance at two AUC cut offs (0.2 and 0.5) indicating more common feature importance with a higher AUC cutoff.

**Suppl. Figure 3:**
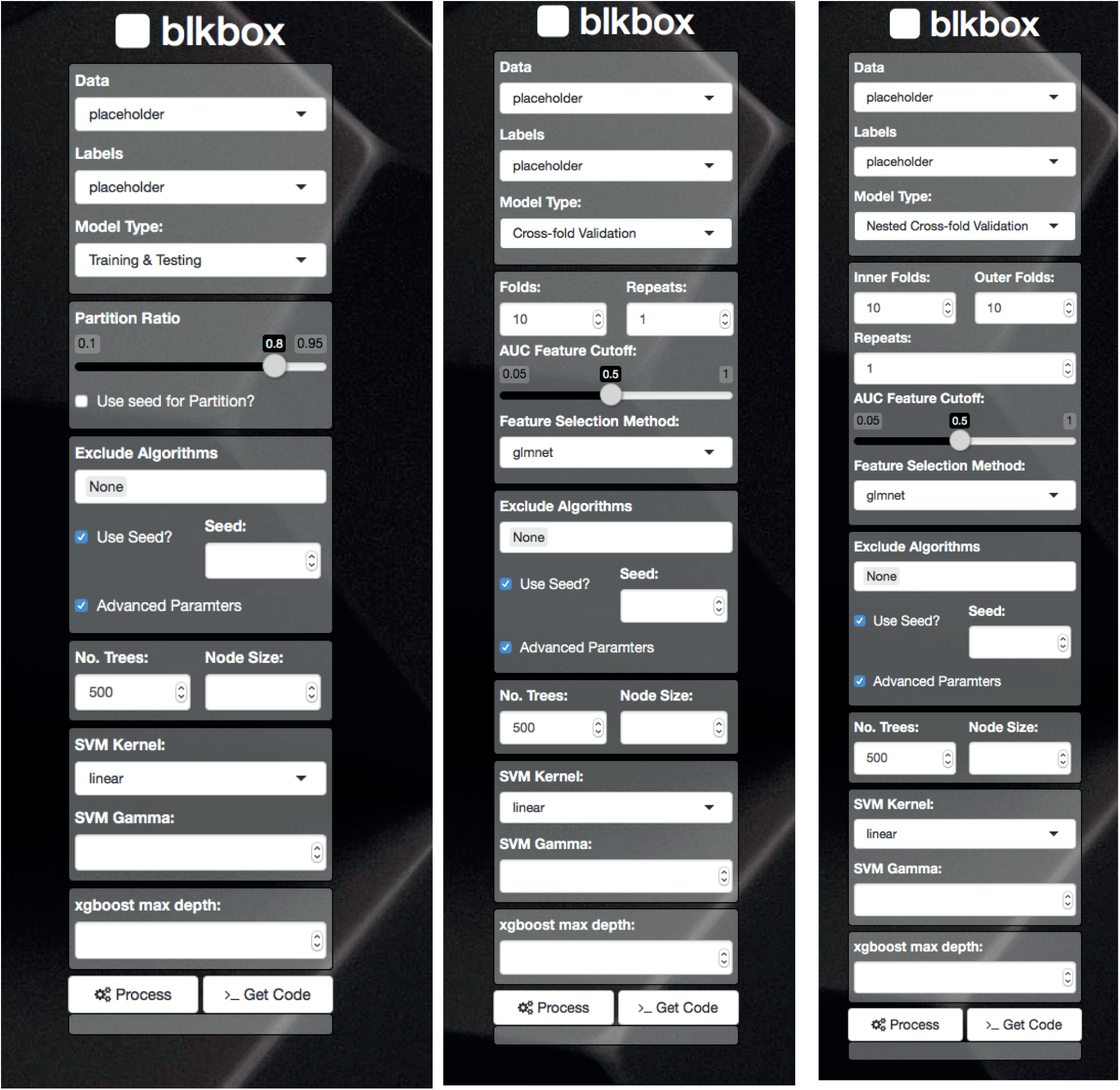
blkbox provides a shiny interface that will enable a “code-free” approach. To begin using the shiny interface run blkboxUI. The interface will allow the submission of a model directly to the current R session, however it will also have the capacity to provide the code used to generate the model. By providing the code as a text output one can easily paste this into any exisitng scripts (e.g. running blkbox on a high performance computing cluster as a submitted job).

**Suppl. Figure 4:**
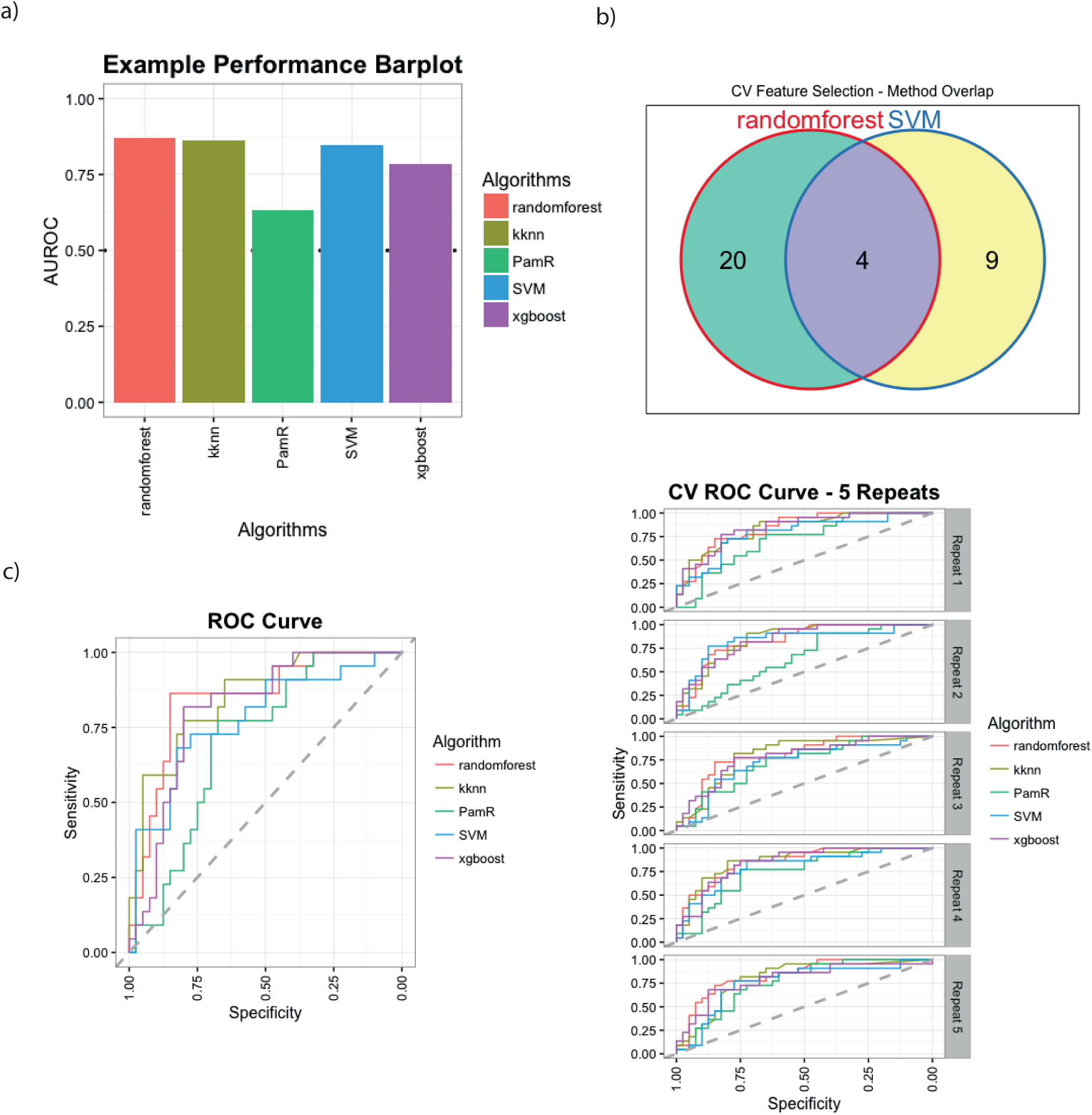
a) Performance measure. blkbox currently supports five different performance metrics; error rate, area under the receiver operating characteristic curve, Matthews correlation coefficient and F-score. Metrics can be applied using the Performance function which will allow one more of the metrics to be specified. cv.plot has the Performance function embedded within it and therefore will not require calling Performance manually. b) Venn Diagrams. When comparing multiple algorithms it can be insightful to visualise the overlap between them, therefore cv.venn and ncv.venn allow comparison of what features were found to be important in each algorithm. cv.venn can be used if more than one Method and a singular AUC were specified when generating a blkboxCV model. cv.venn will compare and intersect the features that survived the AUC cutoff in the specific algorithms (Methods). c) Receiver operating characteristic (ROC) curves are a measure of true positive and false positive rates, the area under this curve is often used as a metric for evaluating model performance. blkbox offers the blkboxROC function that uses the pROC package to calculate the curve and then feeds that to ggplot2 for an aesthetic overhaul. If repeatitions were run with blkboxCV for the models then the plot becomes faceted by repeat number. Similiarly when running blkboxNCV the plot can be generated for holdouts individually or combined, the former is faceted whilst the later is a singular ROC curve.

**Suppl. Figure 5:**
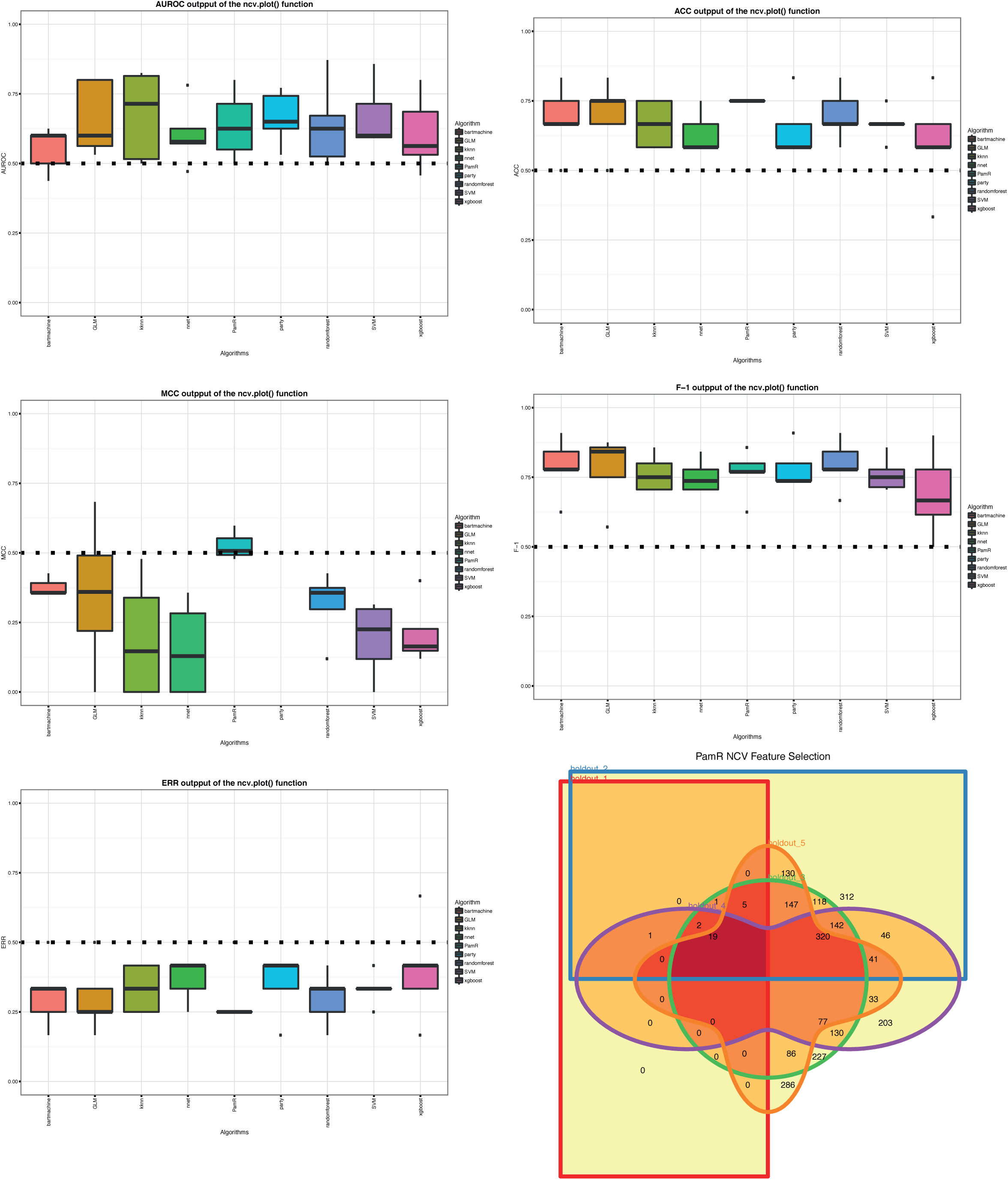
Different metric output of the ncv.plot() and ncv.venn() function. ncv.plot is similar to cv.plot but does not recalculate the performance and uses only the exisiting metrics within a blkboxNCV model. ncv.venn varies from cv.venn in a major way, it doesnt directly compare algorithms but compares the features used in each holdout of blkboxNCV. This can be a measure of feature stability and be informative to adjustments made on the AUC and other parameters.

## Supplementary Methods

### Function description

The standard Blkbox() function allows the user to partition the data on their own and then train a model for various algorithms before testing it on the holdout data partition.

The BlkboxCV() function extends the training and testing function to a cross validation workflow. It uses k-fold cross validation to partition the input data into k subsets, training the algorithms on k-1 subsets and then evaluating on the remaining subset. Once the data is split into k folds and then k models are generated where the holdout fold is rotated through all possibilities. After the successful completion of k-fold cross validation it may be specified to repeat the process n times, if this option is chosen the data will be shuffled with a pre-determined seed to avoid the same folds as in prior iterations. Blkbox writes a table of the classification of each sample in each algorithm facilitating the extraction of any information desired, such as the consensus of one algorithm over all samples. This consensus is the basis for a confusion matrix, which facilitates robust evaluation according to a chosen performance metric (see above).

BlkboxNCV() wraps around both functions mentioned before to determine the success of a feature selected dataset (based on the training set) on the external holdout. This procedure was necessary to implement since we discovered high accuracy even on randomized negative datasets. The high performance on randomized negative data can be facilitated through undiscovered subsets of features providing perfect classification and is supported by the problematic inherent in p>>n datasets (Grate, 2005).

A key component of the BlkboxNCV is the feature selection based on the area under the curve (AUC) of the combined feature importance. A measure of feature importance can be calculated for all of the algorithms except the kNN method. The individual feature importance measure is calculated for every fold of each repeat, averaged and then scaled between 0 and 100. The calculated metric directly represents a feature’s relative importance to resolve classes and facilitates comparison between algorithms; continuing with features of non-zero relative importance dramatically reduces the computational time whilst maintaining accuracy. (AUC over AUROC figure)

The concatenation of all unique non-zero features is a robust feature selection method that maintains every feature of importance. The matrix of relative feature importance can then be weighted by the models average performance and consistently strong features are both excellent biomarker candidates as well as potentially providing mechanistic insights into the basis of the disease.

blkbox implements nine representative high-performance ML algorithms: Random Forests (rf) (Breiman, 2001), Support Vector Machines (SVM) (Cortes and Vapnik, 1995), neural networks (nnet) (Naylor, 1996), k-Nearest Neighbors (kNN) (Hechenbichler and Schliep, 2004), shrunken centroid (pamR) (Tibshirani et al., 2002), conditional inference trees (party) (Hothorn et al., 2012), Bayesian additive regression trees (bart) (Chipman et al., 2010), extreme gradient boosting (xgb) (Chen and Guestrin, 2016) and lasso or elastic-net (glm) (Friedman et al., 2010).

We applied different stringency to the feature selection (AUC cutoffs) in step one to assess the affect of increasing feature size over runtime and accuracy. The number of surviving features as a function of decreasing AUC cutoffs follows a decreasing trend line. The distribution of relative importance remains consistent for each algorithm across the multiple datasets; the relative importance distribution is therefore dependent on the ML algorithm itself. The rate at which the number of surviving features reduces as a function of AUC cutoff remains is consistent for algorithms between datasets.

To measure predictive performance (also referred to as “predictivity”), we used the area under the ROC curve (AUC). The ROC curve is the plot of sensitivity versus 1-specificity for a range of threshold values on the outputs/predictions of the classification algorithms. AUC ranges from 0 to 1, where AUCROC 1 corresponds to a perfectly correct classification of samples, AUCROC 0.5 corresponds to classification by chance, and AUC ROC 0 corresponds to an inverted perfect classification. We chose AUC as the standard predictive performance metric because it is insensitive to unbalanced class prior probabilities, it is computed over the range of sensitivity-specificity tradeoffs at various classifier output thresholds, and it is more discriminative than metrics such as accuracy (proportion of correct classifications), F-measure, precision, etc.

